# Isotropic Three-Dimensional Dual-Color Super-Resolution Microscopy with Metal-Induced Energy Transfer

**DOI:** 10.1101/2021.12.20.473473

**Authors:** Jan Christoph Thiele, Marvin Jungblut, Dominic A. Helmerich, Roman Tsukanov, Anna Chizhik, Alexey I. Chizhik, Martin Schnermann, Markus Sauer, Oleksii Nevskyi, Jörg Enderlein

## Abstract

Over the last two decades, super-resolution microscopy has seen a tremendous development in speed and resolution, but for most of its methods, there exists a remarkable gap between lateral and axial resolution. Similar to conventional optical microscopy, the axial resolution is by a factor three to five worse than the lateral resolution. One recently developed method to close this gap is metal-induced energy transfer (MIET) imaging which achieves an axial resolution down to nanometers. It exploits the distance dependent quenching of fluorescence when a fluorescent molecule is brought close to a metal surface. In the present manuscript, we combine the extreme axial resolution of MIET imaging with the extraordinary lateral resolution of single-molecule localization microscopy, in particular with direct stochastic optical reconstruction microscopy (*d*STORM). This combination allows us to achieve isotropic three-dimensional super-resolution imaging of sub-cellular structures. Moreover, we employed spectral demixing for implementing dualcolor MIET-*d*STORM that allows us to image and co-localize, in three dimensions, two different cellular structures simultaneously.

## Introduction

Super-resolution microscopy has revolutionized optical imaging by extending the limits of spatial resolution by three orders of magnitude down to a few nanometers. The first truly super-resolving microscopy methods were Stimulated Emission Depletion (STED) microscopy^[1]^ and later REversible Saturated OpticaL Fluorescence Transitions (RESOLFT)^[2]^ developed by Stefan Hell and coworkers. This pioneering work spurred the development of another class of super-resolution methods, Single-Molecule Localization Microscopy (SMLM), which is based on the idea that one can localize the center position of an individual fluorescent molecule with much higher accuracy than the width of the molecule’s image (defined by the optical resolution of a microscope). SMLM comprises methods such as Stochastic Optical Reconstruction Microscopy (STORM),^[3]^ PhotoActivatable Localization Microscopy (PALM),^[4]^ Point Accumulation for Imaging in Nanoscale Topography (PAINT)^[5]^ microscopy, its commonly used variant DNA-PAINT,^[6]^ or *direct* STORM (*d*STORM).^[7]^

All of the above mentioned methods provide superb lateral resolution, but cellular structures are of course intrinsically three-dimensional. Thus, several approaches have been developed to extend the super-resolution capabilities to the third dimension. For STED, the use of special phase plates allows for generating stimulated emission intensity distributions with particular resolution enhancement along the optical axis.^[8]^ For SMLM, different techniques have been introduced such as biplane imaging,^[9]^ astigmatic imaging,^[10]^ or various point spread function (PSF) designs such as double-helix PSF,^[11]^ corkscrew PSF,^[12]^ or Tetrapod PSF^[13]^. Recently, clever PSF phase self-modulation has been used for three-dimensional SMLM deep in tissue.^[14]^ However, all these techniques provide an axial resolution that is by a factor 3-5 worse than the achievable lateral resolution, very similar to the resolutions achieved in conventional, diffraction-limited confocal laser scanning microscopy (CLSM).

This gap between lateral and axial resolution was closed by 4π interferometric microscopy techniques that interfere the emission of a molecule detected from two opposite sides with two objectives. This leads to a dramatic improvement in axial resolution as demonstrated by interferometric PALM (iPALM),^[15]^ isoSTED,^[16]^ or whole-cell 4Pi single-molecule switching nanoscopy (W-4PiSMSN).^[17]^ However, these methods are based on macroscopic interferometers that are experimentally very challenging to operate, which prevented their wide distribution and application so far. One of the latest additions to the zoo of 3D SMLM is 3D-MINFLUX.^[18]^ With 3D-MINFLUX, it is possible to localize single molecules with sub-nanometer accuracy by detecting as few as some hundred photons.^[19]^ Moreover, the recently introduced pulsed interleaved MINFLUX (p-MINFLUX) simplifies the experimental setup making it potentially more amenable for wider use.^[20]^ However, the currently existing versions of MINFLUX suffer from low throughput (number of localized molecules per time) and are still technically more complex than almost all SMLM methods that are based on conventional wide-field microscopes.

An attractive alternative to the above mentioned interferometric methods are techniques based on evanescent fields. The first of these approaches uses the exponentially decaying excitation intensity in a total internal reflection fluorescence microscope (TIRFM), where the sample is illuminated from the glass side with a plane wave incident under a high angle above the critical angle of total internal reflection (TIR). That generates an evanescent electromagnetic field on the sample side, so that the excitation intensity that a molecule sees depends on its distance from the surface. By taking several snapshots for excitations under different excitation angles, and thus modulating the exponential decay of the evanescent field intensity, it is possible to calculate distances of molecules (fluorescent structures) from the surface with a few nanometer precision (variable angle TIRFM or vaTIRFM).^[21, 22]^ Alternatively, one can use the evanescent field of fluorescence emission for measuring molecule-surface distance values. One of the first realizations of this idea was super-critical angle fluorescence detection, which uses the fact that the evanescent field of an emitting molecule can couple into propagating light modes on the glass side, which can then be detected with an objective of sufficiently high numerical aperture. This coupling efficiency is again highly distance dependent, due to the evanescent nature of the coupled field. By comparing the intensity of this supercritical emission (named so for its emission angles above the critical TIR angle) with “classical” emission below the critical TIR angle (which does not depend on molecule-surface distance) one can again deduce distance values of single molecules with an accuracy of few nanometers.^[23–25]^

Another technique that exploits the evanescent field of fluorescence emission is Metal-Induced Energy Transfer (MIET).^[26]^ The technique uses the distance-dependent coupling of the evanescent field of a fluorescent emitter to surface plasmons in a thin metallic layer deposited on the surface of the glass cover slide. The resulting energy transfer is extremely distance dependent and leads to a distance-dependent fluorescence lifetime and intensity of the emitter, which can be used to determine molecule-distance values with nanometer accuracy (single-molecule MIET or smMIET),^[27–29]^ despite the unavoidable fluorescence intensity losses due to partial light absorption by the metal film. This is due to the fact that, although the fluorescence brightness of a dye is increasingly reduced the closer the dye comes to the metal surface, its photo-stability increases proportionally, so that the average number of detectable photons from one molecule until photobleaching is nearly independent on dye-metal distance. Due to the broad absorption spectra of metals, the energy transfer from a fluorescent molecule to the metal takes place with high efficiency across the full emission spectrum of a molecule. Meanwhile, MIET imaging was successfully employed for studying various biological questions, for example blood platelet spreading and adhesion,^[30]^ the reorganization of the actin cytoskeleton during epithelial to mesenchymal cell transformation,^[31]^ or the measurement of the inter-bilayer distance of a nuclear envelope.^[32]^ An interesting alternative to a metal film as energy acceptor is graphene, which shows a much steeper lifetime-versus-distance dependence,^[33]^ and which allows for achieving an order-of-magnitude better axial localization accuracy, down to a few Angstrom.^[34–36]^

Thus, a combination of MIET imaging with the high lateral resolution of SMLM could provide isotropic three-dimensional super-resolution imaging of cellular structures. However, SMLM techniques traditionally utilize wide-field imaging while MIET requires precise single molecule lifetime measurements that typically rely on CLSMs. To overcome this problem, we recently introduced CLSM for fluorescence lifetime SMLM (FL-SMLM) imaging.^[37]^ This technique has several advantages in comparison to wide-field SMLM, like a light exposure limited to only the scanned area and optical sectioning that allows imaging deeply into the cell. But most importantly, it provides lifetime-information on a single molecule basis which enables lifetime-based multiplexing within the same spectral window and therefore allows for chromatic aberration-free super-resolution imaging of multiple cellular structures.

In this work, we present a combination of smMIET with *d*STORM, one of the most powerful and widely used SMLM techniques. Our approach combines all the advantages of FL-SMLM with the exquisite axial resolution of MIET imaging. Firstly, we demonstrate MIET-*d*STORM on imaging DNA-labelled polymer beads and surface-immobilized dsDNA-constructs. To show that MIET-*d*STORM can be used for a wide range of biological applications, we imaged microtubules and clathrin coated pits in fixed U2OS and COS-7 cells. Moreover, dual-color MIET allowed for simultaneous imaging of both structures when utilizing spectral demixing dSTORM (sd-*d*STORM).^[38]^

## Results and Discussion

### Validation of MIET-SMLM

In MIET-SMLM, the axial information is encoded in the fluorescence lifetime. To access the single-molecule lifetimes, we preformed FL-SMLM with a custom-built confocal microscope with a fast laser scanner, single-photon detection, and TCSPC electronics (for more details see Figure S1).

For validation of the method, and to check the axial precision of smMIET, we immobilized Alexa Fluor 647-dsDNA-biotin constructs on a gold-coated cover glass topped with a SiO_2_ spacer layer of well-defined thickness. We used the dye Alexa Fluor 647 (AF 467) for labeling which is known for its good performance in dSTORM measurements. Measured TCSPC curves (Figure 1a) and single molecule lifetime histograms (Figure 1b) show the expected lifetime increase with increasing spacer thickness. From the MIET measurements, we deduce that the BSA-neutravidin immobilization layer has a thickness of ~12 nm which is in excellent agreement with literature values.^[39]^ The width of the height distributions (Figure 1d) reflects the surface roughness and axial localization precision. Therefore, the data confirms that the axial localization precision is below 10 nm up to a height of 60 nm.

**Fig. 1.**
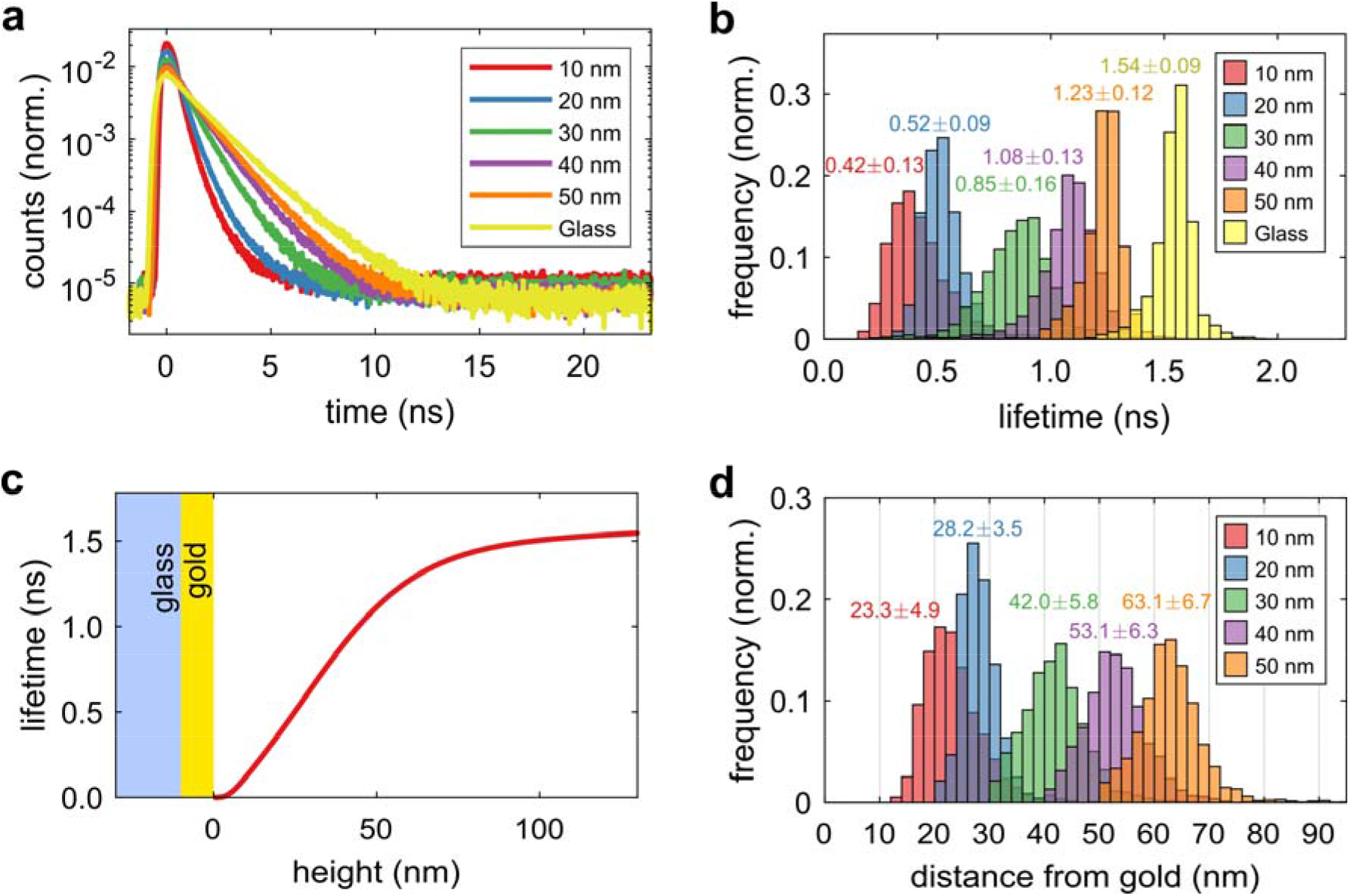
MIET-SMLM validation. **(a)** TCSPC curves for DNA labelled with AF 647 on MIET substrates with different SiO_2_ spacers and on pure glass. **(b)** Single molecule lifetime histograms of DNA labelled with AF 647 on MIET substrates with different SiO_2_ spacers and on pure glass. The lifetime histograms include data from several regions of interest. **(c)** MIET-curve for AF 647 above a MIET substrate with a 10 nm gold layer. **(d**) Histograms of axial positions (height values) of single molecules calculated with the MIET-curve from their measured lifetimes. Averages and standard deviations of lifetime and height values are given next to each peak.

### Imaging biological structures utilizing MIET-SMLM

3D imaging with MIET-SMLM is compatible with biological samples. To demonstrate this, cells were seeded on a cover glass coated with 10 nm of gold and 5 nm of SiO_2_ using standard immunofluorescence sample preparation procedures (see methods for details). The SiO_2_ layer is crucial to protect the gold from the chemically reductive environment during sample preparation and from the thiols in the imaging buffer. Due to their well-defined structure, microtubules are a popular benchmark sample. Therefore, we first imaged α-tubulin in U2OS cells (see Figure 2a) which were chosen due to their planarity. The diffraction limited FLIM image (Figure 2b) already reveals clear lifetime differences along the microtubules but the finer details become only visible in the FL-SMLM reconstruction. For each single molecule, lifetime values were converted to height values to obtain its 3D position. In Figure 3c, a super-resolved reconstruction from 3D localizations, subtle height differences on the order of a microtubule diameter become visible in the network. MIET-SMLM does not compromise the lateral localization precision, which we estimated to be 9.1 nm using a modified Mortensen equation.^[40, 41]^

**Fig. 2.**
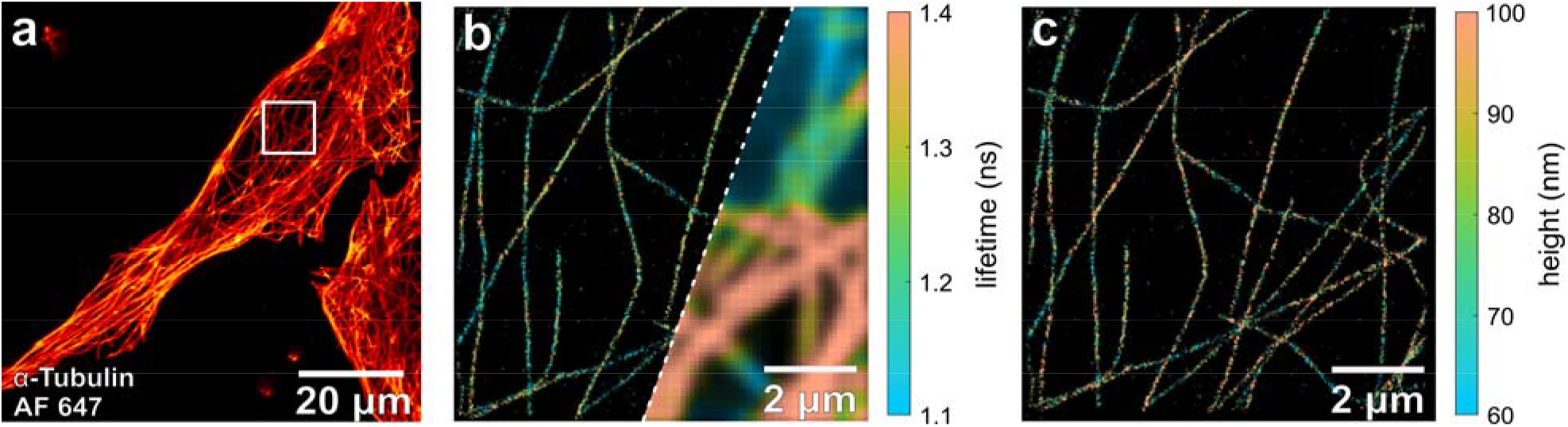
MIET-*d*STORM imaging in cells. **(a)** Confocal laser-scanning image of α-tubulin filaments in U2OS cells labelled with AF 647. **(b)** Confocal FLIM and super-resolved FLIM image of the region-of-interest marked in (a). **(c)** Super-resolved height image of the corresponding region-of-interest.

**Fig. 3.**
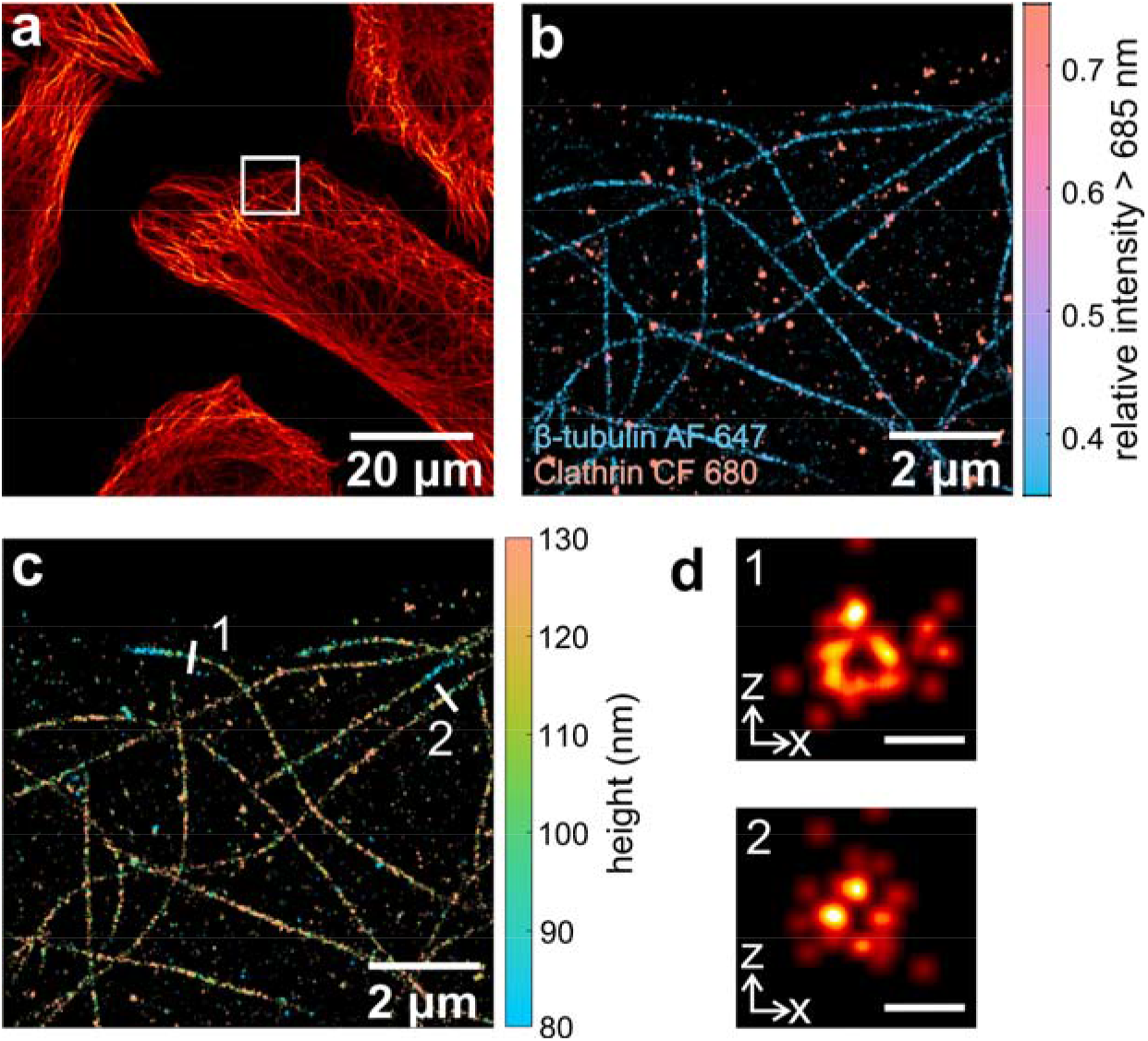
Simultaneous dual-color MIET-*d*STORM imaging in cells. **(a)** Diffraction-limited confocal laser-scanning image of β-tubulin and clathrin in COS-7 cells labelled with AF 647 and CF 680, respectively. **(b)** Sd-*d*STORM image of the region-of-interest marked in (a). **(c)** Three-dimensional MIET-*d*STORM image of the region-of-interest marked in (a), where lifetime values were converted to height values, and both targets are shown together. **(d)** *xz* cross-sections of microtubules *1* and *2* shown in (a). Scale bar is 50 nm.

Employing MIET does not restrict the choice of possible fluorophores. We have performed MIET-SMLM with several types of fluorophores, such as AF 647 and CF 680 for classical *d*STORM imaging, and Cy5b for reductive caging SMLM (see Figure S2).^[42]^ This demonstrates that MIET-SMLM is completely independent of the switching mechanism or measurement conditions. For correctly modeling MIET imaging as required for data evaluation (conversion of lifetime into distance values), exact knowledge of emission spectra, fluorescence quantum yields and fluorescence lifetimes of the used fluorophores (in the absence of any metal quenching) is required. Therefore, we performed lifetime reference measurements on fluorophores far away from the gold-coated cover glass, and we determined absolute values of fluorescence quantum yield of antibody-conjugated fluorophores utilizing a recently developed nanocavity-method (see Table S1).^[43]^

### Simultaneous dual-color MIET-SMLM

The nature of excitation and detection in a CLSM facilitates extension to spectrally-resolved imaging. We implemented dual-color detection by splitting the fluorescence signal, with an additional dichroic mirror, into two separate detection channels, each equipped with a single-photon sensitive detector (for details, see Methods section and Figure S1). With this system, we performed sd-*d*STORM on COS-7 cells with AF 647 labelled α-tubulin and CF 680 labelled clathrin. The spectral and photophysical properties of these two fluorophores make them good candidates for spectral demixing (see Figure S3). In the spectral-resolved reconstruction shown in Figure 3b, it is straightforward to distinguish the two targets α-tubulin and clathrin. The different relative intensities of the two dyes in the two detection channels allows for classification of single molecules with negligible crosstalk and separate reconstruction of the two targets (Figure S4). Spectral-splitting CLSM has the advantage that no channel registration is required and that, due to single-photon counting with almost zero dark counts, the signal-to-noise ratio is excellent. Both aspects are important for achieving highest lateral localization precision, which was estimated to be 9.0 nm for both targets. Moreover, the spectral splitting does not interfere with the lifetime measurement. Measured lifetime values of AF 647 and CF 680 were converted to height values using the corresponding MIET curve for each fluorophore. In Figure 3c, a 3D-*d*STORM image of both targets is presented. Separate super-resolved height images for α-tubulin and clathrin are shown in Figure S4. For both targets, we find structures at height values from below 80 nm to above 130. To highlight the quality of the obtained 3D data, we plotted x-z cross-sections of the microtubules marked in Figure 3d. The hollow structure and the size of the microtubules match theoretical expectations when taking into account that the labelling with secondary antibodies adds an additional distance between the fluorophores and the imaged structures.^[44]^ Our data confirms that MIET-STORM archives high localization precision in all three dimension in complex, biological samples.

### Conclusions

In this work, we presented a new method for 3D super-resolution microscopy. The combination of the high axial precision of MIET imaging with the high lateral resolution of SMLM allows for isotropic single molecule localization in 3D. The achieved axial localization precision is below 10 nm within the first 60 nm from the gold coated cover glass surface. By adding spacer layers or choosing a different substrate, such as graphene,^[45]^ the axial range and sensitivity could be adapted to a given sample. MIET-SMLM is straightforward to implement on commercial CLSMs with TCSPC capability and fast laser scanning. We have demonstrated MIET-SMLM utilizing *d*STORM for imaging cellular structures. Moreover, dual-color MIET imaging via spectral demixing allowed for simultaneous imaging of two different biological structures without compromising resolution. MIET-SMLM could become a powerful tool for multiplexed 3D super-resolution microscopy with exceptionally high isotropic resolution and manifold applications in structural biology.

## Materials and Methods

### Confocal microscope

Fluorescence lifetime measurements were performed on a custom-built confocal setup. Fluorescence excitation was done with a 640 nm 40 MHz pulsed diode laser (PDL 800-B driver with LDH-D-C-640 diode, PicoQuant). After passing through a clean-up filter (MaxDiode 640/8, Semrock), a quarter-wave-plate converted the linearly polarized laser light into circularly polarized light. Subsequently, the laser beam was coupled into a single-mode fiber (PMC-460Si-3.0-NA012-3APC-150-P, Schäfter + Kirchhoff) with a fiber-coupler (60SMS-1-4-RGBV-11-47, Schäfter + Kirchhoff). After the fiber, the output beam was collimated by an air objective (UPlanSApo 10× /0.40 NA, Olympus). An ultra-flat quad-band dichroic mirror (ZT405/488/561/640rpc, Chroma) reflected the excitation light towards the microscope. After passing a laser scanning system (FLIMbee, PicoQuant), the light was sent into the custom side port of the microscope (IX73, Olympus). The three galvo mirrors of the scanning system deflect the beam while preserving the beam position in the back focal plane of the objective (UApo N 100× /1.49 NA oil, Olympus). Sample position is adjusted by using the manual XY stage of the microscope (IX73, Olympus) and a z-piezo stage (Nano-ZL100, MadCityLabs). Fluorescence light was collected by the same objective and de-scanned by the scanning system. An achromatic lens (TTL180-A, Thorlabs) focuses the de-scanned beam onto a pinhole (100 μm P100S, Thorlabs). Backscattered/back-reflected excitation laser light was blocked by a long-pass filter (635 LP Edge Basic, Semrock). After the pinhole, the emission light was collimated by a 100 mm lens. An additional band-pass filter (BrightLine HC 679/41, Semrock) was used for further rejection of scattered excitation light. Finally, the emission light was focused onto a SPAD-detector (SPCM-AQRH, Excelitas) with an achromatic lens (AC254-030-A-ML, Thorlabs).

For sd-*d*STORM, a dichroic mirror (FF685-Di02, Semrock) was used to split the fluorescence signal into two channels, which were focused onto two separate SPAD-detectors. In front of the two detectors, band-pass filters BrightLine HC 679/41 and BrightLine HC 708/75 were placed, respectively (for more details see Figure S1).

Output signals of the photon detectors were recorded with a TCSPC electronics (HydraHarp 400, PicoQuant) that was synchronized by a trigger signal from the excitation laser. Images were acquired with the software SymPhoTime 64 (PicoQuant), which controlled both the TCSPC electronics and the scanner. Typically, samples were scanned with a virtual pixel size of 100 nm, a dwell time of 2.5 μs/pixel, and a TCSPC time resolution of 16 ps.

### MIET imaging

For MIET measurements of COS-7/U2OS cells, samples were prepared on glass coverslips coated with 2 nm titanium, 10 nm gold, 1 nm titanium, and 5 nm silicon dioxide, while for measurements of polymer beads, samples were prepared on glass coverslips coated with 2 nm titanium, 5 nm gold, 1 nm titanium, and 10 nm silicon dioxide. Gold layers were generated by chemical vapor deposition using an electron beam source (Univex 350, Leybold) under high-vacuum conditions (~10^−6^ mbar). A thin silicon dioxide layer of a few nanometers was used for both protecting the gold layer from the thiol buffer and for achieving an optimal distance between sample and gold layer (most sensitive region of MIET curve).

For MIET calibration measurements, we used gold-coated coverslips with SiO_2_ spacers of different thickness on top. The coverslips were rinsed with methanol, and dried using air flow. Four-well silicone inserts (Ibidi 80469, Germany) were attached to a coverslip to form four-well chambers. DNA-fluorophore constructs were immobilized on the surface using biotin-avidin as follows: BSA-biotin (A8549, Sigma-Aldrich) was dissolved and diluted in buffer A (10 mM Tris, 50mM NaCl, pH 8.0) to a concentration of 0.5 mg/mL and added to the chamber and incubated overnight at 4°C. Afterwards, the chamber was flushed with buffer A up to the volume of the chamber for at least 3 times. Neutravidin (31000, Thermo Fisher Scientific) was dissolved and diluted in buffer A to a concentration of 0.5 mg/mL, injected into a chamber and incubated for 5 to 15 min. Then, the neutravidin solution was removed from a chamber by rinsing with buffer A for at least 3 times. Solution with dsDNA-fluorophore at a concentration of 500 pM was added to a chamber and incubated for a few minutes, until sparse coverage of the surface with fluorescent molecules was achieved. The coverage density was controlled visually, and once a desired surface coverage density was reached, the dsDNA leftovers were washed out with B4 buffer (10mM Tris, 1mM EDTA, pH 8.0) including 500 mM NaCl. Imaging was done until all fluorophores photobleached

### Data analysis

Confocal *d*STORM measurements were analyzed with an extended version of the software packed TrackNTrace.^[37, 46]^ From raw scan data, images were generated by combining 10 scans into one frame. When using TrackNTrace, for localization the detection plugin *cross-correlation* with default parameters and the *refinement* plugin *TNT Fitter* with pixel-integrated Gaussian MLE fitting were used. Localizations in adjacent frames with a distance of less than 100 nm were connected to a “track,” and the position was refitted using the sum of all images of the track.

For spectral splitting, localizations were first done on a sum image of both channels. Subsequently, the amplitudes of the Gaussian PSFs were fitted separately in both spectral channels while keeping the PSF size and position fixed.

For lifetime fitting, for each localized molecule a TCSPC histogram was generated by collecting all photons in the corresponding frame with less than 2 σPSF distance from the molecule’s center position. The TCSPC histogram was then fitted with a mono-exponential decay function using a maximum likelihood estimator^[47]^ to determine the lifetime.

Single molecule lifetimes were converted into axial positions using a pre-calculated MIET curve. Localizations were filtered based on PSF size (100 nm < σ_PSF_ < 160 nm), number of photons (> 200), and quality of the lifetime fit (0.9 < Pearson’s χ^2^ < 1.1).

For spectral splitting, molecules were sorted based on the spectral intensity ratio, defined as the intensity in the long wavelength channel divided by the sum of both intensities. Molecules with a ratio below 0.5 were classified as AF 647, molecule above 0.7 as CF 680.

For super-resolution image reconstruction, localizations were reconstructed with a PSF of 15 nm for the large images and 5 nm for the xz-cross sections.

The calibration measurements (Figure 1) were analyzed in a similar fashion to the *d*STORM cell measurements with the following differences: For localization, 100 scans were combined to one frame and molecules not detected in at least two frames were rejected during filtering. For each spacer thickness, the molecule heights were calculated with the corresponding MIET curve. The MIET curve shown in Figure 1c is calculated for a sample without spacer.

The version of TrackNTrace used for this work includes a new plugin for spectral splitting and a data visualizer with added functionalities for MIET, and it is freely available on GitHub (https://github.com/scstein/TrackNTrace).

### Modeling of MIET curves

MIET height-versus-lifetime curves were calculated using published scripts.^[48]^ For this purpose, the geometric structure of the sample (layer composition and thickness values), the numeric aperture of the objective, the emission maximum of the fluorophore, its fluorescence lifetime and its fluorescence quantum yield have to be known. Quantum yield values were adjusted for the actual sample environment by multiplying measured quantum yield values with the ratio of the lifetime measured in the sample to the lifetime measured during quantum yield measurement. In all cases, a random fluorophore orientation was assumed.

### Preparation of dsDNA for surface labelling

The following DNA sequences were used for surface immobilization: the single-stranded DNA (ssDNA *1*) (5^/^ → 3^/^) fluorophore-GCAGCCACAACGTCTATCATCGATT was biotinylated at its 5^/^ end, while its complementary single-stranded DNA (ssDNA *2*) AATCGATGATAGACGTTGTGGCTGC-biotin was labelled with a fluorophore (AF 647) on its 3^/^ end. These two DNA strands were hybridized at high concentration (200 nM) by heating up to 94°C in an annealing buffer for 5 min, and then gradually cooled down to room temperature (30 min). The obtained dsDNA had a length of 25 nucleotides, which ensured its stability on a time scale of several weeks. The construct was designed in such way that the fluorophore faced the surfaces therefore decreasing the linkage errors in single molecule localization. The extra height due to the thickness of the biotin-avidin layer is between 12-16 nm^[29]^ and it was taken into account when estimating the total height above the gold layer.

### dSTORM buffer composition

For conventional *d*STORM imaging (utilizing AF 647 and CF 680), a switching buffer consisting of 50 mM cysteamine in PBS pH 7.4 was used. For reductive single molecule localization microscopy utilizing Cy5b, the following procedure was used: First, the sample was incubated in 0.1% NaBH_4_/ PBS solution for 30 min. Then, it was washed 2-3 times with 0.1% NaBH_4_/PBS and measured in the same 0.1% NaBH_4_/PBS solution. After the measurement, it was washed and stored in PBS.

### Cell culture and antibody labeling

Cell lines were cultured at 37°C in 5% CO_2_ in T25-culture flasks (Thermo Fischer Scientific, #156340). U2OS (human osteosarcoma cell line) and COS-7 (African green monkey kidney fibroblast cell line) were cultivated in Dulbecco’s Modified Eagle Medium (DMEM/F12) with L-glutamine (Sigma, D8062) supplemented with 10% FBS (Sigma-Aldich, F7524) and 100□U/mL penicillin□+□0.1□mg/ml streptomycin (Sigma P4333).

For labeling antibodies with a varying degree of labelling (DOL), an excess of Alexa Fluor 647 NHS-ester (LifeTech, A20106), CF680 NHS-ester (Biotium, #92220), or Cy5B NHS-ester, respectively, was used. The latter was kindly provided by Prof. Dr. Martin Schnermann (National Cancer Institute; Frederick, US-MD).^[49]^ Goat anti-rabbit IgG (IgG-gam, Invitrogen, 31212) and goat anti-mouse IgG (IgG-gar, Sigma-Aldrich, SAB3701063-1) were used as secondary antibodies for staining. For NHS-labelling, 100 μg of antibodies were transferred to 100 mM sodium tetraborate buffer (Fluka, 71999) (pH 9.5) utilizing ZebaTM Spin Desalting Columns 40K MWCO (Thermo Fischer Scientific, #87766) according to the protocol suggested by the manufacturer. Different excesses of NHS-ester dyes were used to achieve different DOLs. For IgG-gar coupled with Alexa Fluor 647, CF680, or Cy5B, an excess of 25x, 15x, and 20x was used to reach a DOL of ~ 8.3. 4.9, and 2.3, respectively. For IgG-gam coupled with Alexa Fluor 647 or CF680, an excess of 25x and 15x was used to reach a DOL of ~ 8.5 and ~ 7.7, respectively. The reaction proceeded for 4 h at RT while protected from light. Labelled antibodies were separated from free dye, washed three times, and reconstituted into PBS (Sigma-Aldrich, D8537-500 ML) using ZebaTM Spin Desalting Columns 40kDa MWCO. Antibody concentration and DOL were determined by UV-vis absorption spectrometry (Jasco V-650).

### Immunostaining

For immunostaining, cells were seeded onto gold-coated coverslips at a concentration of 5·10^4^ cells/coverslip and cultivated overnight at 37°C and 5% CO_2_. For microtubule and clathrin immunostaining, cells were washed with pre-warmed (37°C) PBS, and permeabilized for 2 min with 0.3% glutaraldehyde (GA) + 0.25% Triton X-100 (EMS, 16220 and Thermo Fisher, 28314) in pre-warmed (37°C) cytoskeleton buffer (CB) consisting of 10 mM MES (Sigma-Aldrich, M8250), pH 6.1), 150 mM NaCl (Sigma-Aldrich, 55886), 5 mM EGTA (Sigma-Aldrich, 03777), 5 mM glucose (Sigma-Aldrich, G7021), and 5 mM MgCl_2_ (Sigma-Aldrich, M9272). After permeabilization, cells were fixed with a pre-warmed (37°C) solution of 2% GA in CB for 10 min. After fixation, cells were washed twice with PBS and reduced with 0.1% sodium borohydride (Sigma-Aldrich, 71320) in PBS for 7 min. Cells were again washed three times with PBS before blocking with 5% BSA (Roth, #3737.3) in PBS for 1 h. Subsequently, microtubule samples were incubated with 4 ng/μL rabbit anti-α-tubulin antibody (Abcam, #ab18251) or mouse anti-β-tubulin antibody (Sigma-Aldrich, T8328), and clathrin samples were incubated with 4 ng/μL rabbit anti-clathrin antibody (Abcam, #ab21679) or mouse anti-clathrin antibody (Abcam, #2731) in blocking buffer for 1 h. After primary antibody incubation, cells were washed thrice with 0.1% Tween20 (Thermo Fisher, 28320) in PBS for 15 min. After washing, cells were incubated in blocking buffer with 8 ng/μL of custom labeled secondary antibodies or of commercial IgG-gam-F(ab’)2-Alexa Fluor 647 (DOL ~ 3) (Thermo Fisher, A-21237) for 45 min. After secondary antibody incubation, cells were again washed three times with 0.1% Tween20 in PBS for 15 min. After washing, a post-fix with 4% formaldehyde (Sigma-Aldrich, F8775) in PBS for 10 min was performed followed by three additional washing steps with PBS.

### Fluorescence quantum yield measurements

We used a plasmonic nanocavity and a custom-built scanning confocal microscope for absolute fluorescence quantum yield determination.^[43]^ The cavity mirrors were prepared by chemical vapor deposition of silver on the surface of a clean glass cover slide (bottom mirror) and a plane-convex lens (top mirror) by using a Laybold Univex 350 evaporation machine under high-vacuum conditions (~10^−6^ mbar). The bottom and top mirrors had a thickness of 30 and 60 nm, respectively. The distance between the cavity mirrors was monitored by measuring a white light transmission spectrum using an Andor SR 303i spectrograph and an emCCD camera (Andor iXon DU897 BV). By fitting these spectra with a standard Fresnel model of transmission through a stack of plan-parallel layers, one can determine the precise cavity length (distance between mirrors). Fluorescence lifetime measurements were performed with a custom-built confocal microscope equipped with an objective lens of high numerical aperture (Apo N, 60×/1.49 NA oil immersion, Olympus). A white light laser system (Fianium SC400-4-20) with a tunable filter (AOTFnC-400.650-TN) served as excitation source (λ_exc_ = 640 nm). Collected fluorescence was focused onto the active area of a single photon detection module (MPD series, PDM). Data acquisition was accomplished with a multichannel picosecond event timer (PicoQuant HydraHarp 400). Photon arrival times were histogrammed (bin width of 50 ps) for obtaining fluorescence decay curves. From the obtained lifetime-versus-cavity size curves, absolute values of quantum yields were obtained by fitting an appropriate model.^[43]^

## Acknowledgments

The authors thank Dr. Ingo Gregor for fruitful discussions. J.E., J.C.T., and O.N. are grateful to the European Research Council (ERC) via project “smMIET” (Grant agreement No. 884488) under the European Union’s Horizon 2020 research and innovation program. J.E. is grateful for financial support through Germany’s Excellence Strategy - EXC 2067/1-390729940. M. J. and M.S. acknowledge financial support by the DFG (GRK 2157). M.S. is grateful for financial support by the European Research Council via the ERC Synergy Grant project “ULTRARESOLUTION” (Project No. 951275).

## Author contributions

J.C.T., M.J., D.H., M.S., J.E. and O.N. designed the experiments. J.C.T. and O.N. generated and processed the data. D.H. and M.J. labelled antibodies and prepared cells for dSTORM measurements. R.T. prepared dsDNA for surface labelling. A.C. prepared coverslips for MIET-imaging. A. I. C. performed quantum yield measurements. M. S. performed synthesis of Cy5b dye. J.C.T. wrote the analysis software. J.C.T., M.J., D.H., R.T., M.S., J.E. and O.N wrote and finalized the manuscript.

## Competing interests

The authors declare no conflicts of interest.

## Data and materials availability

The data that support the findings of this study are available from the corresponding author upon reasonable request. The analysis software TrackNTrace is available on Github (https://github.com/scstein/TrackNTrace).

## Notes

https://github.com/scstein/TrackNTrace

### Competing Interest Statement

The authors have declared no competing interest.

### Summary of Updates

Acknowledgment section is changed

## References

[1] S. W. Hell, Science 2007, 316, 1153–1158.

[2] S. W. Hell, S. Jakobs, L. Kastrup, Appl. Phys. A 2003, 77, 859–860.

[3] M. J. Rust, M. Bates, X. Zhuang, Nat. Methods 2006, 3, 793–796.

[4] E. Betzig, G. H. Patterson, R. Sougrat, O. W. Lindwasser, S. Olenych, J. S. Bonifacino, M. W. Davidson, J. Lippincott-Schwartz, H. F. Hess, Science 2006, 313, 1642–1645.

[5] A. Sharonov, R. M. Hochstrasser, Proc. Natl. Acad. Sci. 2006, 103, 18911.

[6] J. Schnitzbauer, M. T. Strauss, T. Schlichthaerle, F. Schueder, R. Jungmann, Nat. Protoc. 2017, 12, 1198–1228.

[7] M. Heilemann, S. van de Linde, M. Schüttpelz, R. Kasper, B. Seefeldt, A. Mukherjee, P. Tinnefeld, M. Sauer, Angew. Chem. Int. Ed. 2008, 47, 6172–6176.

[8] M. Dyba, S. W. Hell, Phys. Rev. Lett. 2002, 88, 163901.

[9] M. F. Juette, T. J. Gould, M. D. Lessard, M. J. Mlodzianoski, B. S. Nagpure, B. T. Bennett, S. T. Hess, J. Bewersdorf, Nat. Methods 2008, 5, 527–529.

[10] B. Huang, W. Wang, M. Bates, X. Zhuang, Science 2008, 319, 810–813.

[11] S. R. P. Pavani, M. A. Thompson, J. S. Biteen, S. J. Lord, N. Liu, R. J. Twieg, R. Piestun, W. E. Moerner, Proc. Natl. Acad. Sci. 2009, 106, 2995.

[12] M. D. Lew, S. F. Lee, M. Badieirostami, W. E. Moerner, Opt. Lett. 2011, 36, 202–204.

[13] Y. Shechtman, L. E. Weiss, A. S. Backer, S. J. Sahl, W. E. Moerner, Nano Lett. 2015, 15, 4194–4199.

[14] P. Bon, J. Linarès-Loyez, M. Feyeux, K. Alessandri, B. Lounis, P. Nassoy, L. Cognet, Nat. Methods 2018, 15, 449–454.

[15] G. Shtengel, J. A. Galbraith, C. G. Galbraith, J. Lippincott-Schwartz, J. M. Gillette, S. Manley, R. Sougrat, C. M. Waterman, P. Kanchanawong, M. W. Davidson, R. D. Fetter, H. F. Hess, Proc. Natl. Acad. Sci. 2009, 106, 3125.

[16] R. Schmidt, C. A. Wurm, A. Punge, A. Egner, S. Jakobs, S. W. Hell, Nano Lett. 2009, 9, 2508–2510.

[17] F. Huang, G. Sirinakis, E. S. Allgeyer, L. K. Schroeder, W. C. Duim, E. B. Kromann, T. Phan, F. E. Rivera-Molina, J. R. Myers, I. Irnov, M. Lessard, Y. Zhang, M. A. Handel, C. Jacobs-Wagner, C. P. Lusk, J. E. Rothman, D. Toomre, M. J. Booth, J. Bewersdorf, Cell 2016, 166, 1028–1040.

[18] F. Balzarotti, Y. Eilers, K. C. Gwosch, A. H. Gynnå, V. Westphal, F. D. Stefani, J. Elf, S. W. Hell, Science 2017, 355, 606.

[19] K. C. Gwosch, J. K. Pape, F. Balzarotti, P. Hoess, J. Ellenberg, J. Ries, S. W. Hell, Nat. Methods 2020, 17, 217–224.

[20] L. A. Masullo, F. Steiner, J. Zähringer, L. F. Lopez, J. Bohlen, L. Richter, F. Cole, P. Tinnefeld, F. D. Stefani, Nano Lett. 2021, 21, 840–846.

[21] M. Cardoso Dos Santos, R. Déturche, C. Vézy, R. Jaffiol, Biophys. J. 2016, 111, 1316–1327.

[22] L. Velas, M. Brameshuber, J. B. Huppa, E. Kurz, M. L. Dustin, P. Zelger, A. Jesacher, G. J. Schütz, Nano Lett. 2021.

[23] C. M. Winterflood, T. Ruckstuhl, D. Verdes, S. Seeger, Phys. Rev. Lett. 2010, 105, 108103.

[24] J. Deschamps, M. Mund, J. Ries, Opt. Express 2014, 22, 29081–29091.

[25] N. Bourg, C. Mayet, G. Dupuis, T. Barroca, P. Bon, S. Lécart, E. Fort, S. Lévêque-Fort, Nat. Photonics 2015, 9, 587–593.

[26] A. I. Chizhik, J. Rother, I. Gregor, A. Janshoff, J. Enderlein, Nat. Photonics 2014, 8, 124–127.

[27] N. Karedla, A. I. Chizhik, I. Gregor, A. M. Chizhik, O. Schulz, J. Enderlein, ChemPhysChem 2014, 15, 705–711.

[28] N. Oleksiievets, J. C. Thiele, A. Weber, I. Gregor, O. Nevskyi, S. Isbaner, R. Tsukanov, J. Enderlein, J. Phys. Chem. A 2020, 124, 3494–3500.

[29] S. Isbaner, N. Karedla, I. Kaminska, D. Ruhlandt, M. Raab, J. Bohlen, A. Chizhik, I. Gregor, P. Tinnefeld, J. Enderlein, R. Tsukanov, Nano Lett. 2018, 18, 2616–2622.

[30] A. Zelená, S. Isbaner, D. Ruhlandt, A. Chizhik, C. Cassini, A. S. Klymchenko, J. Enderlein, A. Chizhik, S. Köster, Nanoscale 2020, 12, 21306–21315.

[31] T. Baronsky, D. Ruhlandt, B. R. Brückner, J. Schäfer, N. Karedla, S. Isbaner, D. Hähnel, I. Gregor, J. Enderlein, A. Janshoff, A. I. Chizhik, Nano Lett. 2017, 17, 3320–3326.

[32] A. M. Chizhik, D. Ruhlandt, J. Pfaff, N. Karedla, A. I. Chizhik, I. Gregor, R. H. Kehlenbach, J. Enderlein, ACS Nano 2017, 11, 11839–11846.

[33] R. J. Moerland, J. P. Hoogenboom, Optica 2016, 3, 112–117.

[34] A. Ghosh, A. I. Chizhik, N. Karedla, J. Enderlein, Nat. Protoc. 2021.

[35] A. Ghosh, A. Sharma, A. I. Chizhik, S. Isbaner, D. Ruhlandt, R. Tsukanov, I. Gregor, N. Karedla, J. Enderlein, Nat. Photonics 2019, 13, 860–865.

[36] I. Kaminska, J. Bohlen, S. Rocchetti, F. Selbach, G. P. Acuna, P. Tinnefeld, Nano Lett. 2019, 19, 4257–4262.

[37] J. C. Thiele, D. A. Helmerich, N. Oleksiievets, R. Tsukanov, E. Butkevich, M. Sauer, O. Nevskyi, J. Enderlein, ACS Nano 2020, 14, 14190–14200.

[38] Z. Zhang, S. J. Kenny, M. Hauser, W. Li, K. Xu, Nat. Methods 2015, 12, 935–938.

[39] K. Spaeth, A. Brecht, G. Gauglitz, J. Colloid Interface Sci. 1997, 196, 128–135.

[40] K. I. Mortensen, L. S. Churchman, J. A. Spudich, H. Flyvbjerg, Nat. Methods 2010, 7, 377–381.

[41] B. Rieger, S. Stallinga, ChemPhysChem 2014, 15, 664–670.

[42] P. Eiring, R. McLaughlin, S. S. Matikonda, Z. Han, L. Grabenhorst, D. A. Helmerich, M. Meub, G. Beliu, M. Luciano, V. Bandi, N. Zijlstra, Z.-D. Shi, S. G. Tarasov, R. Swenson, P. Tinnefeld, V. Glembockyte, T. Cordes, M. Sauer, M. J. Schnermann, Angew. Chem. Int. Ed. 2021, 60, 26685–26693.

[43] A. I. Chizhik, I. Gregor, B. Ernst, J. Enderlein, ChemPhysChem 2013, 14, 505–513.

[44] S. M. Früh, U. Matti, P. R. Spycher, M. Rubini, S. Lickert, T. Schlichthaerle, R. Jungmann, V. Vogel, J. Ries, I. Schoen, ACS Nano 2021, 15, 12161–12170.

[45] I. Kamińska, J. Bohlen, R. Yaadav, P. Schüler, M. Raab, T. Schröder, J. Zähringer, K. Zielonka, S. Krause, P. Tinnefeld, Adv. Mater. 2021, n/a, 2101099.

[46] S. Stein, J. Thiart, Sci. Rep. 2016, 6, 37947.

[47] J. C. Thiele, O. Nevskyi, D. A. Helmerich, M. Sauer, J. Enderlein, Front. Bioinform. 2021, 1, 56.

[48] N. Karedla, A. M. Chizhik, S. C. Stein, D. Ruhlandt, I. Gregor, A. I. Chizhik, J. Enderlein, J. Chem. Phys. 2018, 148, 204201.

[49] P. Eiring, R. McLaughlin, S. S. Matikonda, Z. Han, L. Grabenhorst, D. A. Helmerich, M. Meub, G. Beliu, M. Luciano, V. Bandi, N. Zijlstra, Z.-D. Shi, S. Tarasov, R. Swenson, P. Tinnefeld, V. Glembockyte, T. Cordes, M. Sauer, M. Schnermann, Angew. Chem. Int. Ed. 2021, n/a.

