## Supplementary Materials for "Isotropic Three-Dimensional Dual-Color Super-Resolution Microscopy with Metal-Induced Energy Transfer"

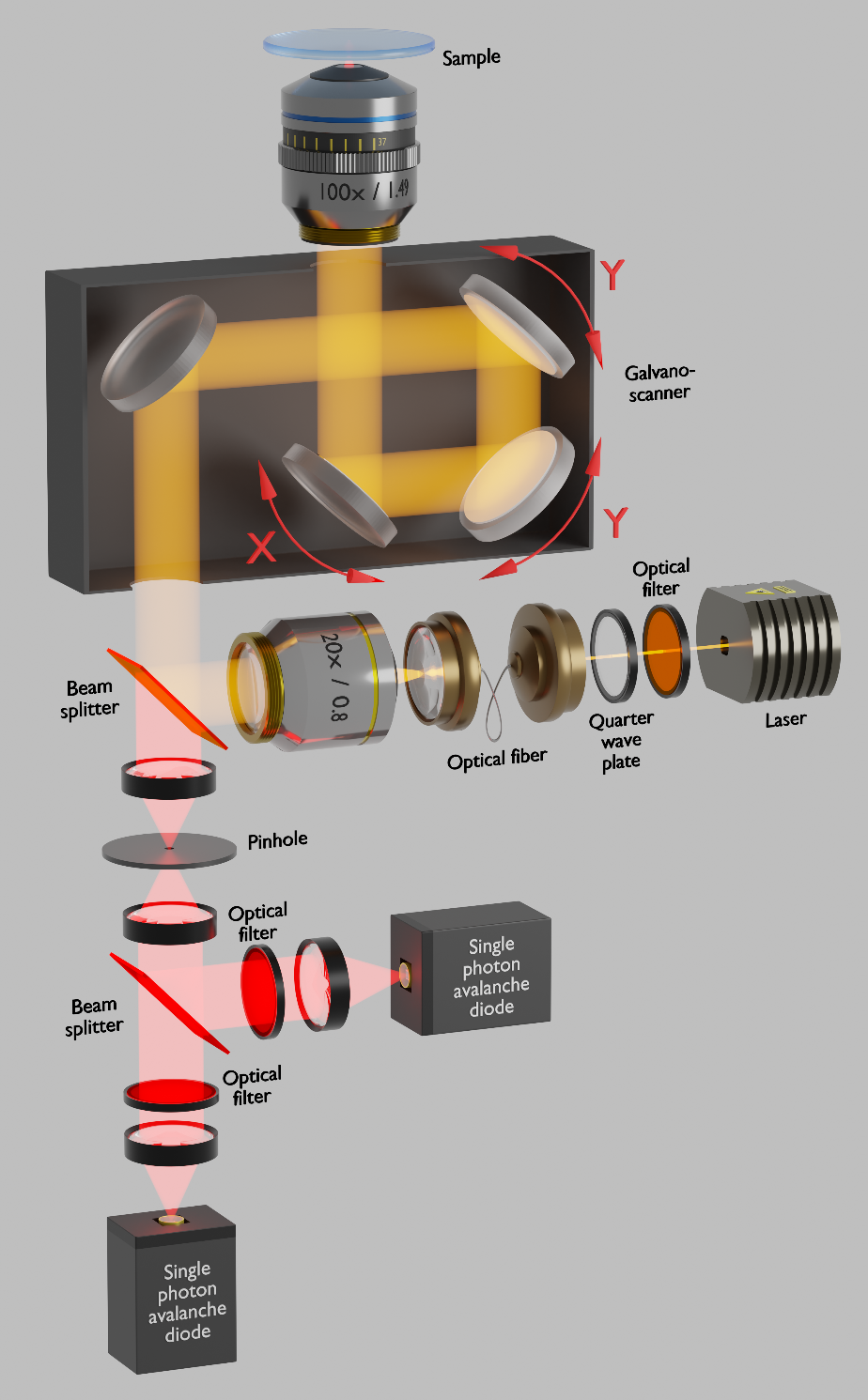

**Fig. S1. Scheme of the MIET-SMLM setup.** The pulsed 640 nm linearly polarized excitation light is converted to circular polarization with a quarter wave plate, passes through a single mode fiber, and is reflected by a non-polarizing beam splitter into a galvo laser scanner and finally focused by the objective into the sample. The collected fluorescence emission from the sample is descanned, passes the beam splitter, and is focused on the pinhole and then on a single-photon detector. For simultaneous dual-color imaging, an additional beam splitter is used. Optical filters (long and band pass filters) are used to block scattered excitation light.

| **Sample** | **Φ** | **τ, ns** |
| --- | --- | --- |
| ssDNA-Alexa647 | 0.28 | 1.36 |
| lgG-gar-Alexa647 | sticks to the surface | |
| lgG-gar-CF680 | 0.21 | 1.25 |
| lgG-gam-CF680 | 0.22 | 1.26 |
| lgG-gar-Cy5B | 0.38 | 1.94 |

**Table S1.	Fluorescence quantum yield and free-space fluorescence lifetime
measurements.**

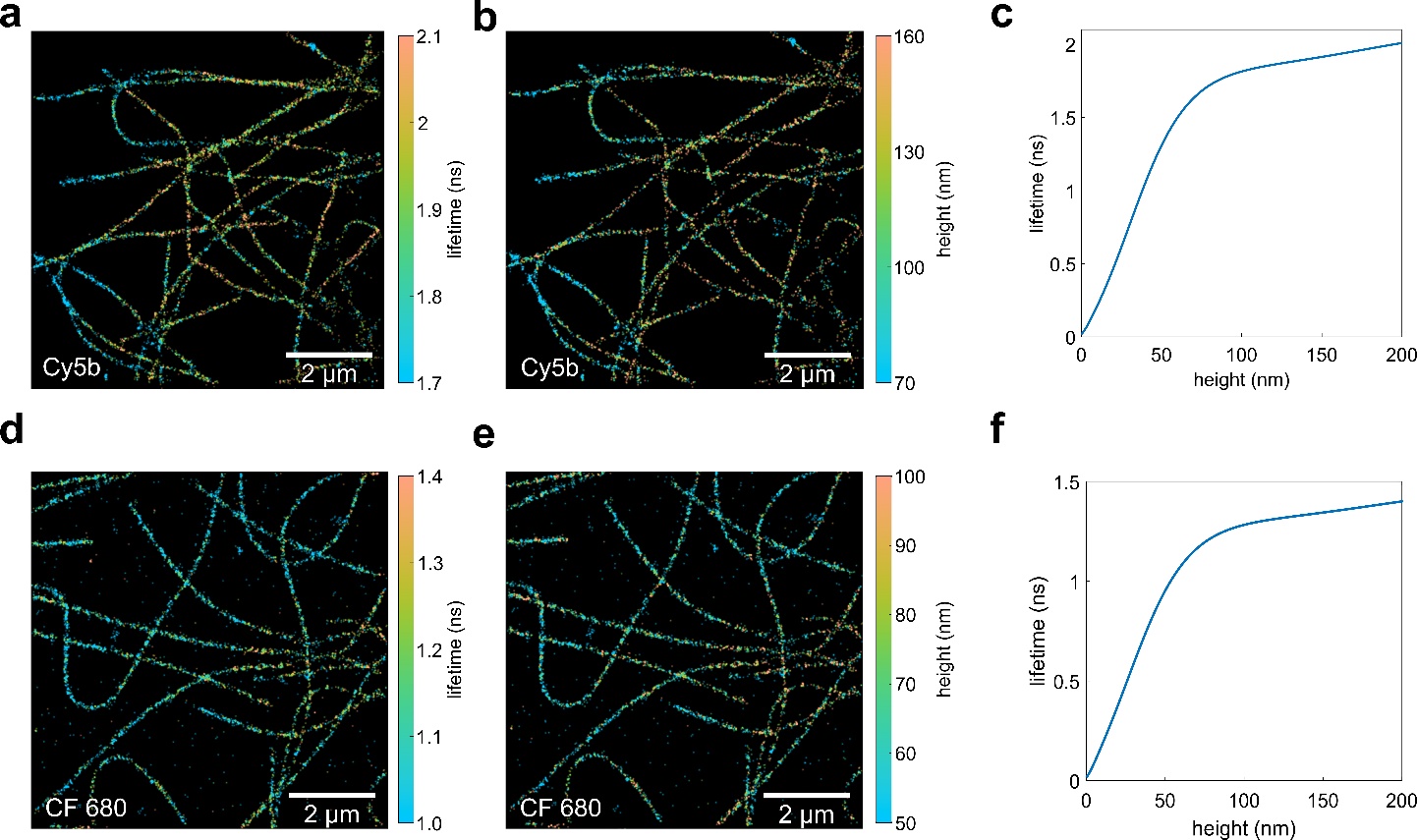

**Fig. S2. MIET-*d*STORM imaging of Cy5b and CF 680 labelled tubulin in U2OS cells.
(a,d)** Lifetime-*d*STORM image of microtubules labelled with Cy5b/CF680. **(b,e)** Corresponding MIET-*d*STORM images, where lifetime values are converted to corresponding height values.
**(c,f)** MIET-curves used for the height determination in Cy5b and CF680 samples, respectively.

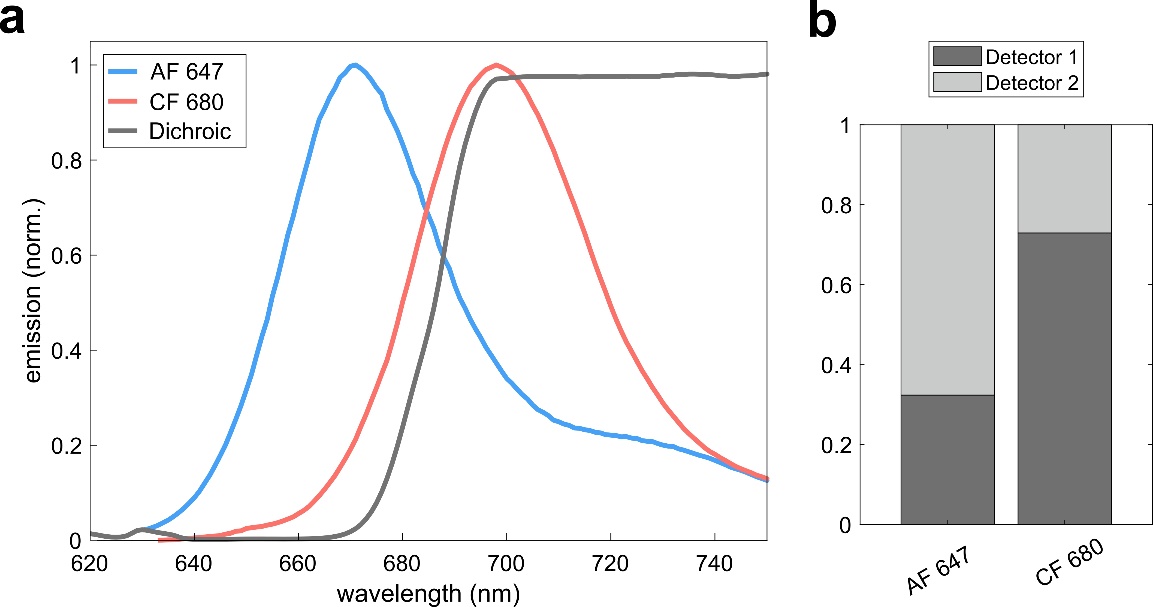

**Fig. S3. Spectral splitting. (a)** Normalized emission spectra of AF 647, CF 680 and transmission of FF685-Di02 dichroic mirror used for spectral splitting. **(b)** Expected intensity ratio between detector 1 and 2 (AF 647 and CF 680).

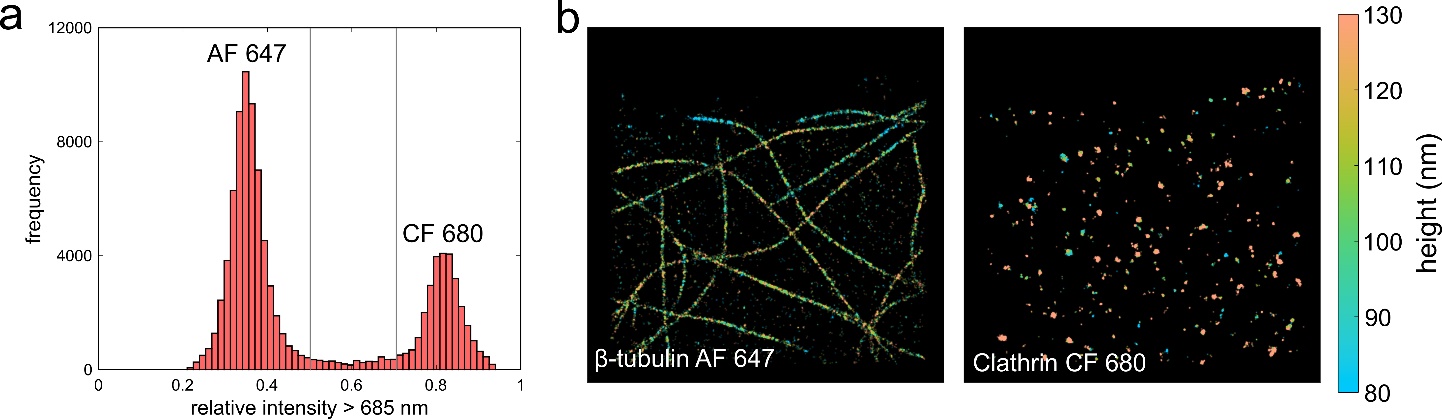

**Fig. S4. Spectrally demixing.** MIET-*d*STORM dual-color imaging of AF 647 labelled β-tubulin and CF 680 labelled clathrin in U2OS cells as presented in Fig. 3. (a) Single molecule histogram of the spectral intensity ratio. (b) Corresponding super-resolved FLIM images for both targets after spectral splitting. The thresholds were a ratio of less than 0.5 for AF 647 and above 0.7
for CF 680.
